# Adipose mTORC2 is essential for arborization of sensory neurons in white adipose tissue and whole-body energy homeostasis

**DOI:** 10.1101/2022.03.21.485116

**Authors:** Irina C Frei, Diana Weissenberger, Danilo Ritz, Wolf Heusermann, Marco Colombi, Mitsugu Shimobayashi, Michael N Hall

## Abstract

Adipose tissue, via sympathetic and sensory neurons, communicates with the central nervous system (CNS) to mediate energy homeostasis. In contrast to the sympathetic nervous system, the morphology, role and regulation of the sensory nervous system in adipose tissue is poorly characterized. Taking advantage of recent progress in whole-mount three-dimensional imaging of adipose tissue, we identified a neuronal network of calcitonin gene-related protein (CGRP)-positive sensory neurons in white adipose tissue (WAT). Furthermore, we show that adipose mammalian target of rapamycin complex 2 (mTORC2), a major component of the insulin signaling pathway, mediates sensory innervation in WAT. Based on visualization of neuronal networks, mTORC2-deficient WAT displayed reduced arborization of (CGRP)-positive sensory neurons, while sympathetic neurons were unaffected. This selective loss of sensory innervation followed reduced expression of growth-associated protein 43 (GAP43) in CGRP-positive sensory neurons. Finally, we found that loss of sensory innervation in WAT correlated with systemic insulin resistance. Our findings suggest that adipose mTORC2 is necessary for sensory innervation in WAT which likely contributes to WAT-to-CNS communication.

## Introduction

White adipose tissue (WAT) stores or releases energy and thereby contributes to whole-body energy homeostasis [1]. Impaired WAT function, often the result of obesity, causes systemic effects such as hyperinsulinemia and insulin resistance [2, 3], indicating that the metabolic state of WAT impacts other tissues. With the increasing prevalence of obesity and type II diabetes, it is important to understand how WAT communicates with distal tissues to control whole-body energy homeostasis.

It is well established that WAT communicates with other tissues, including the brain, by secreting adipokines such as leptin and adiponectin [4, 5]. WAT also interacts with the central nervous system (CNS) via sympathetic and sensory neurons [6-8]. The sympathetic nervous system transmits signals from CNS to WAT. For example, fasting triggers sympathetic neurons to release the neurotransmitter norepinephrine (NE) in WAT. NE then stimulates adipocytes to release free fatty acids that provide energy for other tissues [9]. Compared to the sympathetic nervous system, sensory neurons in WAT are less well studied. Detection of the sensory neuron markers calcitonin gene-related peptide (CGRP), substance P and advillin first suggested the presence of sensory neurons in WAT [10-13]. Anterograde tracing using herpes simplex virus subsequently confirmed the transmission of sensory information from WAT to CNS [14]. Further studies identified the adipokine leptin and bioactive free fatty acids as potential sensory stimuli in WAT [15, 16]. Nevertheless, the morphology, role and regulation of the sensory nervous system in WAT remain poorly understood.

The mammalian target of rapamycin complex 2 (mTORC2) is a multiprotein serine/threonine kinase. It is composed of four core components including the kinase subunit mTOR and the mTORC2-specific subunit rapamycin-insensitive companion of mTOR (RICTOR) [17-19]. Studies of adipose-specific *Rictor* knockout mice revealed that loss of adipose mTORC2 causes reduced glucose uptake and impaired lipid handling in adipocytes [20-25]. Furthermore, loss of adipose mTORC2 non-cell-autonomously causes hyperinsulinemia and systemic insulin resistance [20, 21, 23]. These data suggest that adipose mTORC2 is a critical regulator of systemic energy homeostasis. However, the mechanisms by which adipose mTORC2 mediates communication with other tissues are poorly understood.

We used an inducible adipose-specific *Rictor* knockout (iAdRiKO) mouse to study how adipose mTORC2 mediates WAT communication with other tissues. Phosphoproteomic analysis of inguinal WAT (iWAT) from iAdRiKO mice revealed acute changes in phosphorylation of proteins associated with neurons. To study the effect of adipose mTORC2 on neurons, we visualized sympathetic and sensory neuronal networks in iWAT by performing whole-mount imaging. We discovered that arborization of CGRP-positive sensory neurons, but not tyrosine hydroxylase (TH)-positive sympathetic neurons, was selectively diminished in WAT upon loss of adipose mTORC2. Furthermore, expression of growth-associated protein 43 (GAP43) in CGRP-positive sensory neurons was reduced prior to loss of arborization. We conclude that adipose mTORC2 is required for arborization of sensory neurons in WAT, thereby mediating adipocyte-to-CNS communication.

## Results

### Loss of adipose mTORC2 acutely impairs whole-body energy metabolism

Mice lacking adipose mTORC2 from birth display a defect in whole-body energy homeostasis characterized by hyperinsulinemia and systemic insulin resistance [20, 21, 23]. Since adipose tissue is not fully developed until six weeks of age, this could be due to a developmental defect. To determine whether and, if so, how soon loss of mTORC2 in mature adipocytes causes systemic effects, we performed a longitudinal study with a tamoxifen-inducible *Rictor* knockout (iAdRiKO; *Rictor*^*fl/fl*^, *Adipoq promoter-*CreER^T2^) mouse (**Figure 1A**). Six week old mice were treated with tamoxifen daily for five days and analyzed three days, two weeks, and four weeks after the last tamoxifen treatment (**Figure 1A**). *Cre*-negative mice treated with tamoxifen served as controls. As expected, expression of RICTOR and phosphorylation of the mTORC2 target AKT (AKT-pS473) were downregulated in WAT of iAdRiKO mice at all three post-tamoxifen timepoints (**Figure S1A**). In liver and muscle, RICTOR expression and AKT phosphorylation were not affected (**Figure S1B**). Loss of mTORC2 in mature adipocytes reduced WAT mass approximately 30% compared to control mice (**Figures 1B and S1C**), while brown adipose tissue (BAT) mass was unchanged (**Figure S1D**).

**Figure 1.**
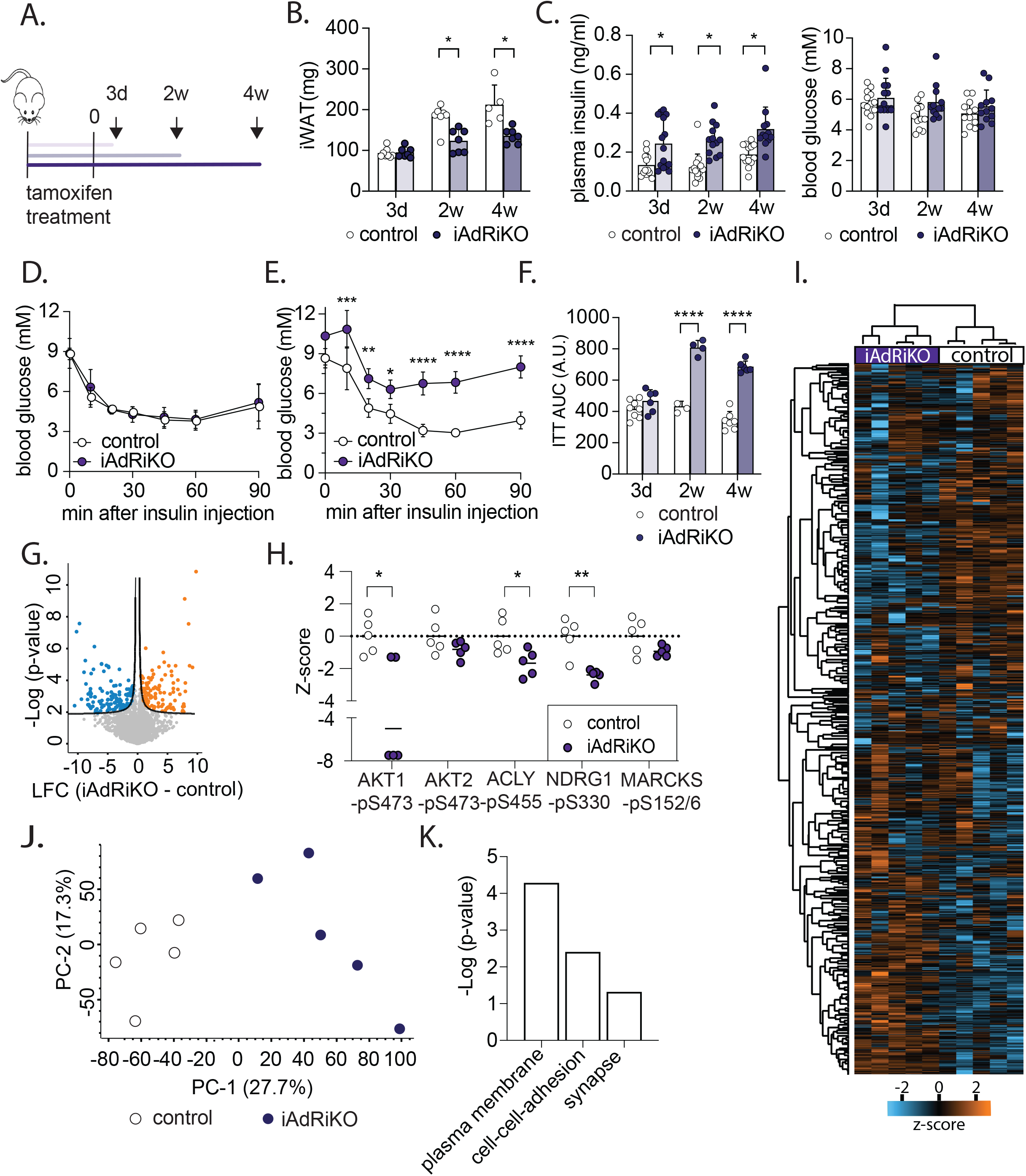
Loss of adipose mTORC2 acutely impairs whole-body energy homeostasis. (A) Experimental design of the longitudinal studies. (B) Tissue weight of inguinal iWAT (iWAT) in *ad libitum*-fed control and iAdRiKO mice three days (n= 8;7), two (n=7), and four weeks (n= 5;7) after tamoxifen treatment. Student’s t-test, *p<0.05. (C) Plasma insulin and blood glucose levels of control and iAdRiKO mice after 16 hours starvation three days (n=13;15), two (15;13), and four weeks (13;12) after tamoxifen treatment. Student’s t-test, *p<0.05. (D) Insulin tolerance test (ITT) on control and iAdRiKO mice three days after tamoxifen treatment (n= 5;8). (E) ITT on control and iAdRiKO mice four weeks after tamoxifen treatment (n=5;6). 2-way ANOVA, *p<0.05, **p<0.01, ***p<0.001, **** p<0.0001. (F) Area under the curve (AUC) for ITT on control and iAdRiKO mice three days (n= 5;8), two (n= 4), and four weeks (n= 5;6) after tamoxifen treatment. Student’s t-test, **** p<0.0001. (G) Volcano-plot displaying the comparison of phosphosites derived from iWAT of control and iAdRiKO mice three days after tamoxifen treatment (n= 5). LFC= Log_2_ fold change. (H) Z-scores of mTORC2-regulated phosphosites in iWAT three days after tamoxifen treatment (n= 5). Student’s t-test, *p<0.05, **p<0.01. (I) Unsupervised hierarchical clustering using Euclidian distance of the phosphoproteome data set (n=5). (J) Principal component analysis (PCA) for iWAT-derived phosphoproteome of control and iAdRiKO mice three days after tamoxifen treatment (n=5). (A) Pathway enrichment analysis of phosphoproteins altered in iWAT of iAdRiKO mice compared to control mice analyzed and visualized by the Database for Annotation, Visualization and Integrated Discovery (DAVID).

Next, we characterized the metabolic state of iAdRiKO and control mice. Fasting insulin levels were acutely increased in iAdRiKO mice and remained significantly elevated while blood glucose levels remained normal (**Figure 1C**). These results suggest that elevated insulin compensates for reduced insulin sensitivity upon loss of mTORC2, as observed previously in mice lacking adipose mTORC2 at birth [20, 23]. In agreement with this suggestion, loss of mTORC2 reduced glucose uptake in WAT (**Figures S1E-F**). Furthermore, iAdRiKO mice become insulin resistant, as determined by an insulin tolerance test (**Figures 1D-F**). Severe insulin resistance was detected in iAdRiKO mice at two and four weeks (**Figures 1E-F and S1G**). Further investigation revealed mild insulin resistance in iAdRiKO mice at five days after tamoxifen treatment (**Figures S1H-I**), indicating a gradual development of insulin resistance within two weeks after loss of adipose mTORC2 (**Figures 1F and S1I**). Taken together, these longitudinal studies revealed rapid development of hyperinsulinemia followed by insulin resistance upon loss of adipose mTORC2. These results highlight the importance of adipose mTORC2 for whole-body energy homeostasis.

### Loss of mTORC2 alters proteins associated with synapse formation in WAT

To gain better understanding of the role of adipose mTORC2 in whole-body energy homeostasis, we performed proteomic and phosphoproteomic analyses on iWAT obtained from iAdRiKO and control mice three days after tamoxifen treatment. This time point allowed us to study the acute effects of reduced mTORC2 activity (**Figure S1A**), before development of systemic insulin resistance (**Figures 1D-F**). The proteomic analysis identified and compared 6’082 proteins in iWAT of iAdRiKO and control mice. Only one protein was significantly altered in AdRiKO mice three days after tamoxifen treatment (40S ribosomal protein S24; p < 0.01; |log_2_(iAdRiKO/control)| > 0.6) (**Figure S2A; Supplementary Table 1A**). In contrast, the phosphoproteomic analysis detected and quantified 10’443 phosphorylated sites of which 319 and 249 were significantly up- and down-regulated (p<0.01), respectively, upon loss of adipose mTORC2 (**Figure 1G; Supplementary Table 1B**). As expected, mTORC2 downstream readouts AKT1-pS473, ACLY-pS455 [26], NDRG1-pS330 [27] and MARCKS-pS152/156 [28] were downregulated in iWAT of iAdRiKO mice compared to control iWAT (**Figure 1H**). Next, we examined whether loss of mTORC2 in adipocytes elicits a molecular signature. Indeed, unsupervised hierarchical clustering analysis (**Figure 1I**) and principal component analysis (PCA) (**Figure 1J**) of the phosphoproteomic data identified two distinct clusters corresponding to iAdRiKO mice and control mice.

To identify molecular processes associated with loss of mTORC2 signaling, we performed pathway enrichment analysis. Proteins whose phosphorylation was significantly altered were associated with the plasma membrane and cell-cell adhesion (**Figure 1K**). This observation is in agreement with previous studies showing that many mTORC2 substrates are localized to the plasma membrane [29, 30]. Unexpectedly, the pathway enrichment analysis also revealed changes in proteins associated with the nervous system, in particular synapse-associated proteins (**Figure 1K**). To investigate the possibility of ectopic *Adipoq promoter-* CreER^T2^ expression and thus mTORC2 ablation in neurons, we examined adiponectin expression in neurons in iWAT. We found no evidence for adiponectin expression in nerve bundles innervating iWAT (**Figure S2B**). This finding is supported by large-scale single-cell RNA sequence data performed in dorsal root ganglion of mice [31], where iWAT-innervating sensory neurons originate. Thus, changes in phosphorylation of synapse-associated proteins are unlikely due to ectopic ablation of mTORC2 in neurons. Based on these findings, we hypothesize that loss of adipose mTORC2 impacts the neuronal network in iWAT. Furthermore, our findings suggest that the acute effects of mTORC2 loss are due to changes in protein phosphorylation rather than expression, as expected for loss of a kinase.

### Loss of adipose mTORC2 reduces arborization of sensory neuron

To investigate the consequences of altered phosphorylation of synaptic proteins, as observed upon acute loss of adipose mTORC2, we examined neuronal innervation in iWAT four weeks after tamoxifen treatment. We first examined the sympathetic nervous system by visualizing TH-positive neurons in iWAT, using a modified whole-mount volumetric imaging protocol [32, 33]. We combined the tissue preparation protocol described by Chi et *al*. (2018a) with a water-based clearing method [34] to enhance structure preservation and antibody combability. As previously described [33, 35], we detected a dense network of TH-positive sympathetic neurons in nerve bundles that branched into smaller fibers running along blood vessels and into the parenchyma (**Figure S3A**). However, we found no change in abundance or morphology of TH-positive sympathetic neurons in iWAT of iAdRiKO mice, compared to controls (**Figure 2A**). In agreement with this result, immunoblot analyses showed no change in TH expression in iWAT of iAdRiKO mice (**Figure 2B**). Next, we analyzed the activity of sympathetic neurons in iWAT. TH-positive neurons release NE to stimulate β-adrenergic signaling in adipocytes, which promotes phosphorylation of hormone sensitive lipase (HSL) at S563 and S660 [36, 37]. Immunoblot analyses revealed no difference in phosphorylation of S563 or S660 (**Figure 2B**), indicating similar sympathetic activity in iAdRiKO and control mice. Taken together, these findings suggest that the morphology and activity of the sympathetic nervous system are normal in iWAT of iAdRiKO mice.

**Figure 2.**
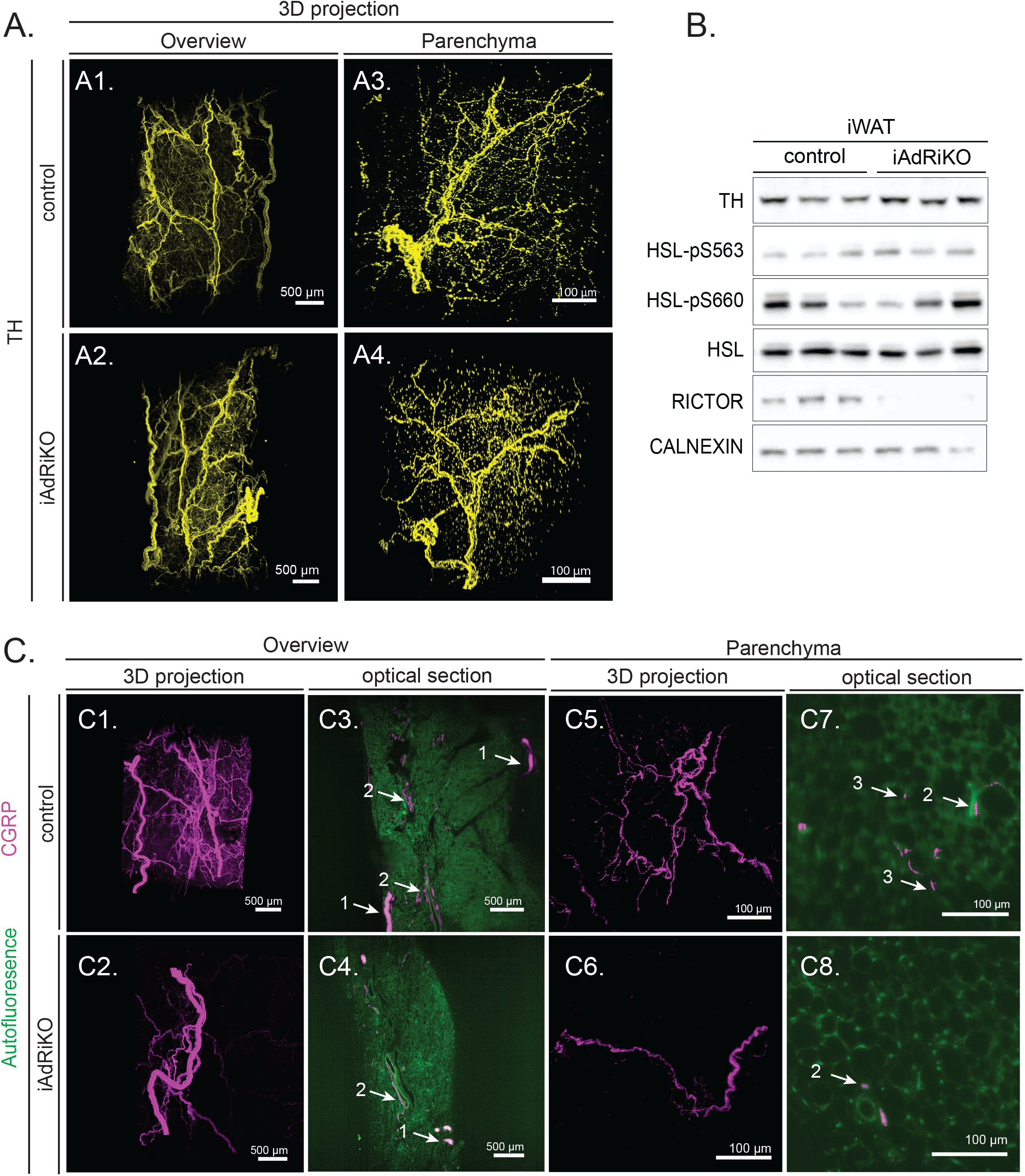
Sensory but not sympathetic innervation is altered upon loss of adipose mTORC2. (A) 2D representatives of a 3D reconstruction of inguinal WAT (iWAT) four weeks after tamoxifen treatment immunostained with tyrosine hydroxylase (TH; yellow). (A1-2) Low magnification projection of sympathetic neuronal network in control and iAdRiKO mice (N=4;5). Scale bar= 500 µm. (A3-4) High magnification projection of sympathetic neurons in iWAT parenchyma of control mice and iAdRiKO (N=19;10). Scale bar= 100 µm. (B) Immunoblot analysis of iWAT from control and iAdRiKO mice four weeks after tamoxifen treatment. Hormone-sensitive lipase (HSL). (n=6;6). (C) 2D representatives of a 3D reconstruction of iWAT four weeks after tamoxifen treatment immunostained with calcitonin gene-related peptide (CGRP; magenta). (C1-2) Low magnification projection of sensory neuronal network in control and iAdRiKO mice (N=12;19). Scale bar= 500 µm. (C3-4) Low magnification cross section of sensory neuronal network in control and iAdRiKO mice (N=12;19). Nerve bundle (1), innervation along blood vessel (2), tissue autofluorescence (green). Scale bar= 500 µm. (C5-6) High magnification projection of sensory neurons in iWAT parenchyma of control mice and iAdRiKO (N=16;11). Scale bar= 100 µm. (C7-8) High magnification cross section of neurons in control and iAdRiKO mice (N=16;11). Innervation along blood vessel (2), parenchymal innervation (3), tissue autofluorescence (green). Scale bar= 100 µm.

We next examined the sensory nervous system in iWAT. Since the three-dimensional sensory neuronal network in iWAT was not previously described, we first examined CGRP-positive sensory neurons in control mice. Similar to TH-positive neurons, CGRP-positive sensory neurons innervated iWAT via large nerve bundles (**Figure 2C**). From the large nerve bundles, smaller nerve bundles branched off and ran along blood vessels into the tissue (**Figure 2C**). Using high magnification imaging, we detected single nerve fibers branching out into the parenchyma where sensory neurons were in close contact with adipocytes (**Figure 2C**).

Next, we analyzed the sensory neuronal network in iWAT of iAdRiKO mice. CGRP-positive sensory neurons were present in large and small nerve bundles similar to control mice (**Figures 2C and S3B)**. Importantly, unlike control mice, we did not detect CGRP-positive neurons in the parenchyma, indicating loss of arborization of sensory neurons in iWAT of iAdRiKO mice (**Figures 2C and S3B, Movies 1 and 2**). Loss of CGRP in iWAT lacking mTORC2 was also observed by an enzyme-linked immunosorbent assay (ELISA) (**Figure S3C**). To further investigate the loss of CGRP-positive sensory neurons in iWAT, we co-stained and visualized TH- and CGRP-positive neurons in whole iWAT depots collected from iAdRiKO and control mice. We discovered that TH- and CGRP-positive neurons were present in the same larger and smaller nerve bundles and ran alongside each other into the tissue (**Figures 3A and S4A-B, Movie 3**). However, the TH-positive sympathetic neuronal network was much denser compared to the sensory neuronal network, particularly in the parenchyma. Confirming the above observations, sensory neurons were not detected in the parenchyma of iAdRiKO iWAT, while sympathetic neurons were detected (**Figures 3B-C and S4C, Movies 4 and 5**). Taken together, these findings suggest that adipose mTORC2 is required for arborization of CGRP-positive sensory neurons, but not of sympathetic neurons, in iWAT.

**Figure 3.**
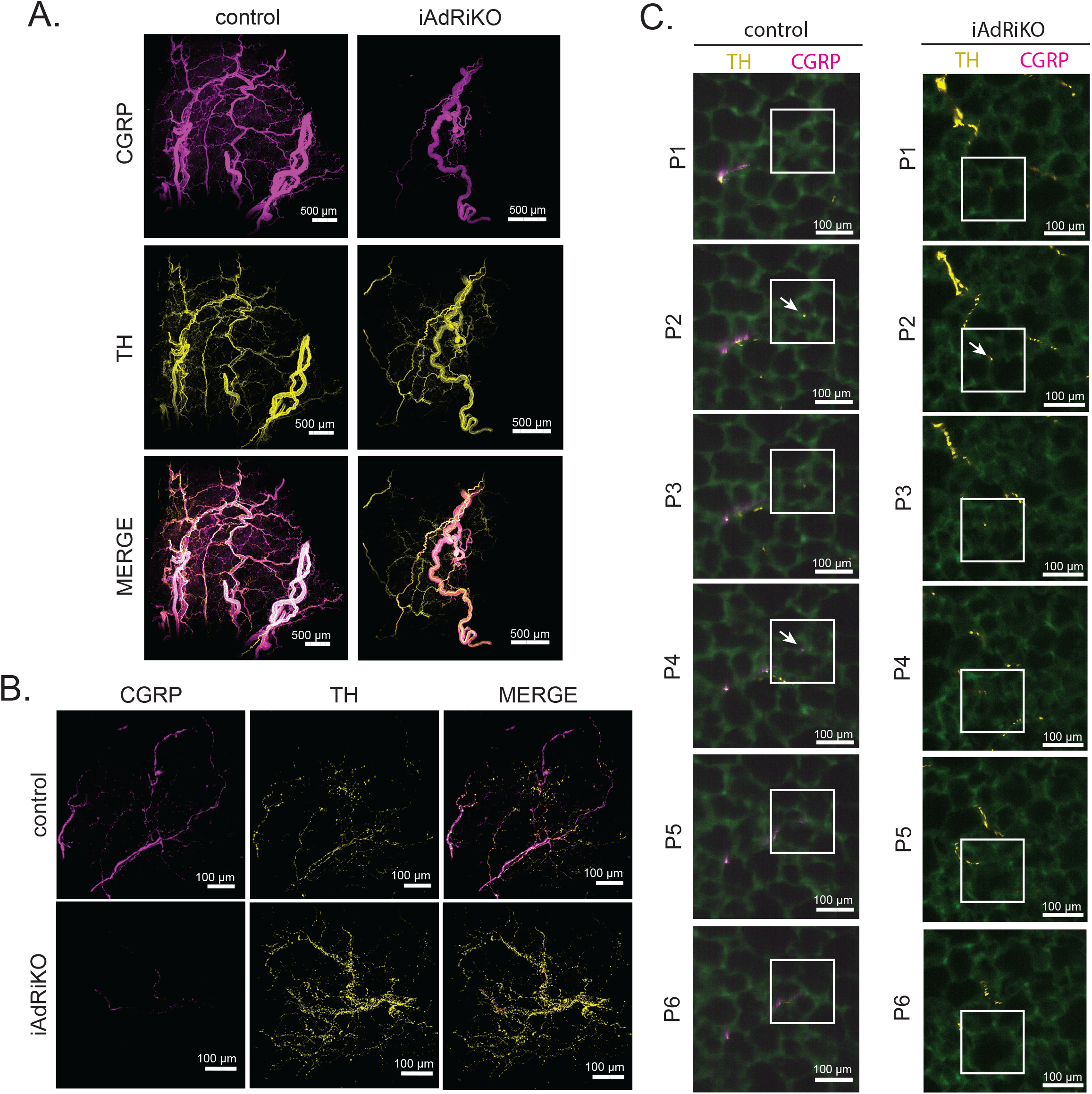
Loss of adipose mTORC2 reduces arborization of sensory neurons. (A) Low magnification projection of inguinal WAT (iWAT) four weeks after tamoxifen treatment co-immunostained with tyrosine hydroxylase (TH, yellow) and calcitonin gene-related peptide (CGRP, magenta) (N=15;18). Scale bar= 500 µm. (B) High magnification projection of iWAT four weeks after tamoxifen treatment co-immunostained with TH (yellow) and CGRP (magenta) (N=21;22). Scale bar= 500 µm. (C) Subsequent sections (P1-P6; 6.48/6.28 µm interval, respectively) of control and iAdRiKO iWAT immunostained with TH (yellow) and CGRP (magenta) illustrating single nerve fibers innervating the periphery. Region of interest: Nerve ending and potential synapses (arrows). Tissue autofluorescence= green. Scale bar= 100 µm.

We noted that the intensity of CGRP staining was slightly reduced in the main nerve bundles of iAdRiKO iWAT (**Figure 2C**). To confirm this observation, we performed co-staining of TH and CGRP in paraffine sectioned iWAT (**Figure S4D**). The number of CGRP-positive single fibers was similar in nerve bundles of iAdRiKO and control mice. However, the intensity of CGRP per fiber was reduced 40% in iAdRiKO iWAT (**Figures S4D-E**). Consistent with the whole-mount imaging (**Figure 2A**), we detected no change in TH intensity in iWAT of iAdRiKO mice (**Figures S4D and S4F)**. The reduction of CGRP in single nerve fibers, in addition to loss of arborization, may indicate a loss in sensory activity.

### GAP43 expression is downregulated in CGRP-positive neurons upon loss of adipose mTORC2

We next investigated how loss of adipose mTORC2 reduces arborization of CGRP-positive neurons in iWAT. We examined neuronal growth-associated protein 43 (GAP43), a protein that promotes neuronal growth and plasticity in the CNS [38]. GAP43 was previously shown to be downregulated in iWAT of *ob/ob* mice, a mouse model for type II diabetes [39]. Immunoblot analysis of total and phosphorylated (pS41) GAP43 showed a strong decrease in GAP43 expression in iWAT lacking mTORC2, at two and four weeks after tamoxifen treatment (**Figures 4A-B**). Since WAT is a heterogenous tissue, we next examined whether GAP43 is expressed only in neurons. We determined GAP43 expression in surgically denervated iWAT, observing that GAP43 expression was indeed lost upon denervation (**Figure 4C**). These data suggest that GAP43 is expressed only in neurons and thus the loss of GAP43 observed in iWAT of iAdRiKO is due to loss in the nervous system.

**Figure 4.**
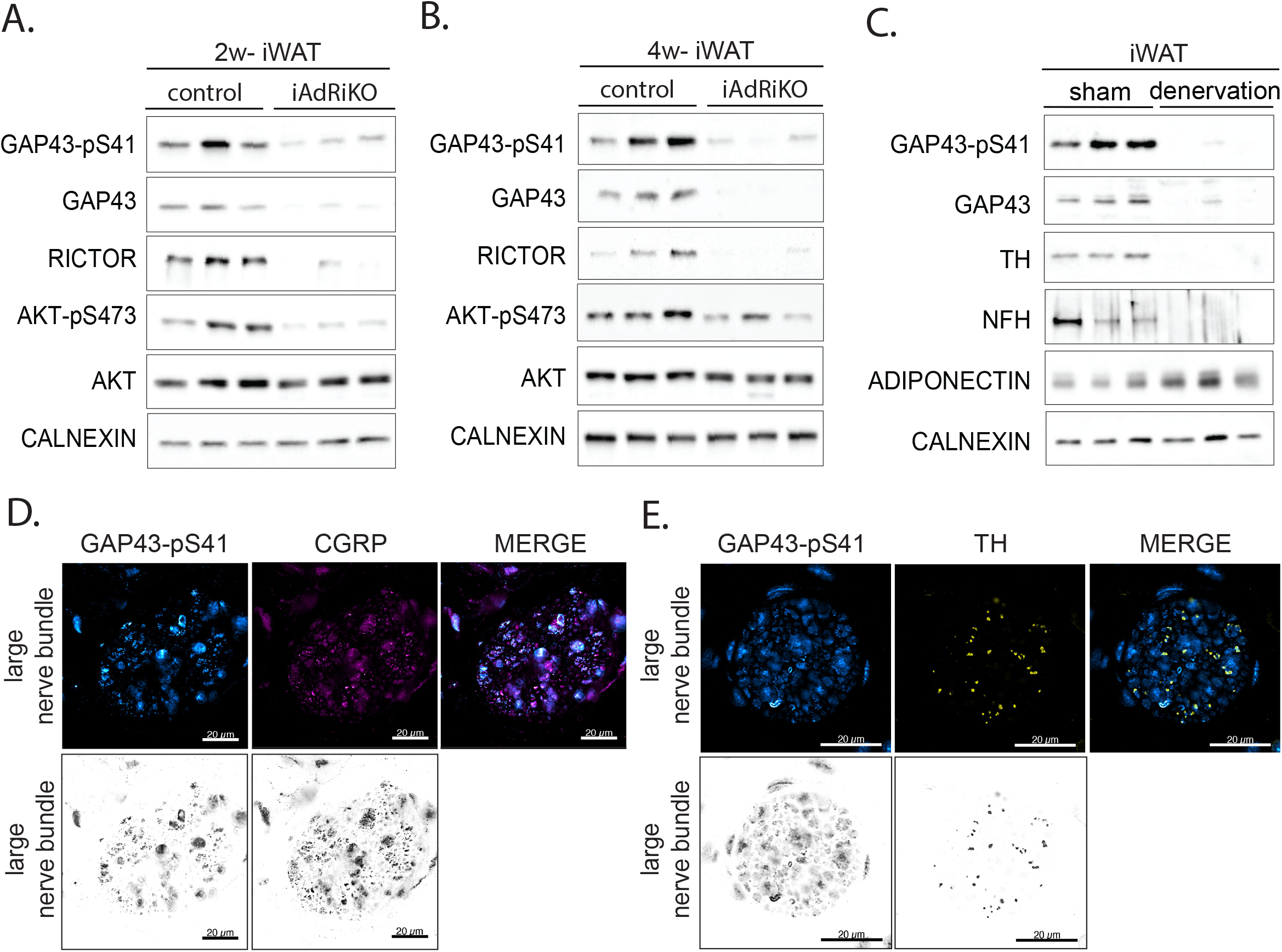
GAP43 expression is downregulated in CGRP-positive neurons upon loss of adipose mTORC2. (A) Immunoblot analysis of inguinal WAT (iWAT) tissue from control and iAdRiKO mice two weeks after tamoxifen treatment. (n=6;6). (B) Immunoblot analysis of iWAT tissue from control and iAdRiKO mice four weeks after tamoxifen treatment. (n=6;6). (C) Immunoblot analysis of surgically denervated iWAT depot (denervation) compared to iWAT depot from sham-operated mice (sham). Neurofilament heavy polypeptide (NFH). (n=5;5). (D) Representative image of a large nerve bundle in iWAT of control mice immunostained with growth-associated protein 43 (GAP43)-pS41 and calcitonin gene-related peptide (CGRP). (N=11;9). (E) Representative image of a large nerve bundle in iWAT of control mice immunostained with GAP43-pS41 and tyrosine hydroxylase (TH). (N=19;11).

Previous reports showed that GAP43 can be expressed in both sensory and sympathetic neurons [40, 41]. To address which class of neurons express GAP43 in iWAT, we visualized GAP43 in TH- or CGRP-positive neurons in large nerve bundles in sectioned iWAT. We note that we visualized GAP43 with antibody against phosphorylated GAP43 because antibody against total GAP43 did not serve as an immunostaining reagent. We found that the GAP43-pS41 signal correlated with CGRP-positive fibers, but not with TH-positive fibers, indicating that GAP43 is expressed specifically in sensory neurons in iWAT (**Figures 4D-E**). Considering the established role of GAP43 in neuronal growth [38], our data suggest that loss of adipose mTORC2 reduces arborization of CGRP-positive neurons possibly due to loss of neuronal GAP43 expression.

## Discussion

In this study, we provide insights on the morphology, role and regulation of the sensory nervous system in WAT. Taking advantage of recent progress in whole-mount three-dimensional imaging, we show that sensory neurons are present in central nerve bundles, along blood vessels and in the parenchyma of WAT. Furthermore, we discovered that loss of mTORC2 signaling in adipocytes decreases arborization of sensory neurons without affecting sympathetic innervation or activity. Our findings suggest that adipose mTORC2 promotes growth (arborization) of CGRP-positive sensory neurons in WAT, thereby supporting adipocyte-to-CNS communication.

The morphology of the sensory nervous system in WAT has not been described previously. We show that CGRP-positive sensory neurons, like TH-positive sympathetic neurons, innervate the parenchyma of WAT. We note that the sensory nervous system forms a less dense network compared to the sympathetic nervous system. Nevertheless, sensory neurons appear to contact adipocytes (**Figure 3C**). The nature of these connections is still unknown, since sensory neurons may form stable synapses with adipocytes or terminate as free nerve endings. However, the close proximity of adipocytes and sensory neurons provides morphological evidence for adipocyte-to-CNS communication via the sensory nervous system.

The role of sensory neurons in WAT is poorly understood. It has been proposed that sensory neurons may directly communicate the metabolic state of WAT to CNS [14-16]. Indeed, leptin and species of free fatty acids have been identified as potential sensory stimuli acting in WAT [15, 16]. Our data provide additional evidence for this hypothesis since arborization of sensory neurons is lost upon ablation of adipose mTORC2, a critical regulator of energy uptake and storage in WAT [23, 25]. In addition, the observed reduction of sensory innervation in WAT lacking mTORC2 coincides with hyperinsulinemia and insulin resistance. Accordingly, CGRP-positive neurons in iWAT may impact whole body energy homeostasis via the CNS, although ablation of CGRP-positive neurons specifically in BAT had no impact on whole-body energy homeostasis [42]. Further studies are needed to elucidate the potential role of WAT-derived sensory neurons in controlling energy homeostasis.

How does adipose mTORC2 regulate growth of CGRP-positive sensory neurons in WAT? We observed that loss of mTORC2 correlates with loss of the neuronal growth promoting protein GAP43. An intriguing possibility is that adipose mTORC2 promotes GAP43 expression in sensory neurons. Since GAP43 is expressed only in sensory neurons (**Figures 4D-E**), this hypothesis explains why CGRP-positive neurons, but not TH-positive neurons, are affected in iWAT of iAdRiKO mice. GAP43 is found at the presynaptic membrane and at neuronal growth cones [38]. Our phosphoproteomic analysis revealed that phosphorylation of both membrane-associated proteins and cell-cell adhesion proteins was altered upon loss of adipose mTORC2 (**Figure 1K**). Thus, loss of mTORC2 in adipocytes may disrupt the postsynaptic membrane, thereby destabilizing synapses and affecting pre-synaptic GAP43 expression. Alternatively, CGRP-positive sensory neurons may require a secreted stimulus from adipocytes to undergo arborization. It has been suggested that sensory neurons respond to secreted free fatty acids produced by lipolysis or *de novo* lipogenesis in adipocytes [16, 43]. Since mTORC2 promotes *de novo* lipogenesis [23, 44], loss of mTORC2 may decrease production and thus release of free fatty acids by adipocytes which may in turn decrease arborization of sensory neurons. Further studies are required to uncover the molecular mechanism underlying adipose mTORC2-mediated regulation of CGRP-positive sensory neurons in iWAT.

Loss of sensory neurons has been observed in diabetic patients, a condition known as diabetic-induced neuropathy [45]. However, the underlying mechanism(s) is poorly understood. Blaszkiewicz et al. have reported that GAP43-positive neurons are lost in iWAT in genetically obese (*ob/ob*) mice [39]. Since a decrease in mTORC2 has previously been shown in omental WAT of obese patients [46], it will also be of interest to investigate a possible correlation between loss of CGRP-positive neurons and reduced mTORC2 activity in WAT as a potential mechanism of diabetes-induced neuropathy.

## Acknowledgements

We thank Sara Merino and Kai Schleicher (Imaging Core Facility, Biozentrum, Basel) for technical support, and Stefan Offermanns for providing *Adipoq promoter-*CreER^T2^ mice. We also thank Christoph Handschin, Bernhard Thorens, Markus Rüegg and Frédéric Preitner for helpful discussions. We acknowledge support from the Swiss National Science Foundation (project 179569 to MNH and 161510 to MS), The Louis Jeantet Foundation (MNH), and the Canton of Basel (MNH).

## Author contributions

ICF, MS, and MNH designed the project and the experiments. ICF, DW and MS conducted the experiments with contributions from DR (proteomics), MC (proteomic analysis) and WH (clearing method). ICF, MS and MNH wrote the manuscript.

## Materials and Methods

### Mice

As described before, the iAdRiKO mouse line was generated by breeding *Rictorfl/fl* mice with *adipoq-CreERT2* mice, provided by Stefan Offermanns (Max Planck Institute for Heart and Lung Research [MPI-HLR], Bad Nauheim, Germany) [46, 47]. For experiments, *Rictorfl/fl adipoq-CreERT2* mice were bred with *Rictorfl/fl* mice to generate iAdRiKO and control littermates. *Rictor* knockout was induced at 6 weeks of age. For that, iAdRiKO and control littermates were treated with i.p. injection daily with 1 mg/mouse tamoxifen (Sigma-Aldrich) resuspended in corn oil for consecutive 5 days. Mice were housed at 22 °C in a regulation compliant facility under a 12-hour light/12-hour dark cycle with unlimited access to water and normal diet (unless stated otherwise). Mice were sacrificed early in the morning and tissues were collected and weighed. All mouse experiments were performed according to federal guidelines for animal experimentation and were approved by the Kantonales Veterinäramt of the Kanton Basel-Stadt under the cantonal license 2602 and 2975.

### 2-Deoxyglucose uptake assay

Mice were fasted for five hours, then injected i.p. with Humalog insulin (Lilly; 0.75 U/kg body weight), followed 10 minutes later with an injection of 2-deoxyglucose (Sigma-Aldrich; 32.8 μg/g body weight). Tissues were collected 20 minutes after administration of 2-deoxyglucose. Tissues were lysed in 10 mM Tris-HCL, pH 8.0, by boiling for 15 minutes. 2-Deoxyglucose-6-phosphate (2DGP) was measured using a Glucose Uptake-Glo Assay Kit (Promega) following the manufacturer’s instructions.

### Insulin tolerance test

Mice were fasted for 5 hours and blood samples were collected to determine blood glucose levels. Humalog insulin was given i.p. (Lilly; 0.75 U/kg body weight), and blood glucose levels were monitored by an Accu-Chek blood glucose meter for 90 minutes.

### Immunoblots

Tissue were homogenized in lysis buffer containing 100 mM Tris-HCl pH 7.5, 2 mM EDTA, 2 mM EGTA, 150 mM NaCl, 1% Triton X-100, cOmplete inhibitor cocktail (Roche) and PhosSTOP (Roche). Protein concentration was determined by Bradford assay. Equal amounts of protein were separated by SDS-PAGE and transferred onto nitrocellulose membranes (GE Healthcare). The nitrocellulose membranes were blocked with 5% BSA in TBST (TBS containing 0.1% Tween20) and incubated overnight in primary antibody diluted in TBST containing 5% BSA. Primary antibodies used were RICTOR (1:1000; Cell signaling; Cat#2140), AKT (1:1000; Cell signaling, Cat#4685), AKT-pS473 (1:1000; Cell signaling, Cat#9271), CALNEXIN (1:1000, Enzo, Cat#ADI-SPA-860-F), GAP43 (1:1000, Cell signaling, Cat#8945), GAP43-pS41 (1:1000, R&D Systems, Cat#PPS006), Tyrosine hydroxylase (1:500, Millipore, Cat#AB1542), HSL (1:2000, Cell signaling, Cat#4107), HSL-pS660 (1:1000, Cell signaling, Cat#4126), HSL-pS563 (1:1000, Cell signaling, Cat#4139). The primary antibody was washed several times with TBST and then incubated in secondary antibody in TBST containing 5% milk powder (w/v). Secondary antibodies used were mouse anti-rabbit (1:10’000, Jackson, 211-032-171) and rabbit anti-sheep (1:10’000, Invitrogen, 81-8620).

### Sample preparation for proteomics and phosphoproteomics

Tissues were pulverized and homogenized in lysis buffer containing 100 mM Tris-HCl pH7.5, 2 mM EDTA, 2 mM EGTA, 150 mM NaCl, 1% Triton X-100, cOmplete inhibitor cocktail (Roche) and PhosSTOP (Roche). Samples were lysed by polytron followed by ultrasonication. Lysates was further cleared from debris and excessive amount of lipids by centrifugation (2×14’000g, 10min). Proteins were precipitated by trichloroacetic acid (Sigma), washed with cold acetone and resuspended in buffer containing 1.6 M urea, 0.1 M ammonium bicarbonate and 5 mM TCEP. Proteins were alkylated with 10 mM chloroacetamide and digested with sequencing-grade modified trypsin (enzyme/protein ratio 1:50) overnight. After acidification with 5% TFA, peptides were desalted using C18 reverse-phase spin columns (Macrospin, Harvard Apparatus) according to the manufacturer’s instructions, dried under vacuum and stored at −20 °C until further use.

For TMT-labelling, 25 μg of peptides per sample were labeled with isobaric tandem mass tags (TMT10plex, Thermo Fisher Scientific) as described previously [48]. In brief, peptides were resuspended in 20 μl labeling buffer (2 M urea, 0.2 M HEPES, pH 8.3) and a peptide calibration mixture consisting of six digested standard proteins mixed in different amounts was spiked into each sample before TMT reagents were added, followed by a 1 h incubation at 25 °C shaking at 500 rpm. To quench the labelling reaction, aqueous 1.5 M hydroxylamine solution was added and samples were incubated for another 5 min at 25 °C shaking at 500 rpm followed by pooling of all samples. The pH of the sample pool was increased to 11.9 by adding 1 M phosphate buffer (pH 12) and incubated for 20 min at 25 °C shaking at 500 rpm to remove TMT labels linked to peptide hydroxyl groups. Subsequently, the reaction was stopped by adding 2 M hydrochloric acid until a pH < 2 was reached. Finally, peptide samples were further acidified using 5% TFA, desalted using Sep-Pak Vac 1cc (50 mg) C18 cartridges (Waters) according to the manufacturer’s instructions and dried under vacuum.

### Proteomics

TMT-labeled peptides were fractionated by high-pH reversed phase separation using a XBridge Peptide BEH C18 column (3,5 µm, 130 Å, 1 mm × 150 mm, Waters) on an Agilent 1260 Infinity HPLC system. Peptides were loaded on column in buffer A (20 mM ammonium formate in water, pH 10) and eluted using a two-step linear gradient from 2% to 10% in 5 minutes and then to 50% buffer B (20 mM ammonium formate in 90% acetonitrile, pH 10) over 55 minutes at a flow rate of 42 µl/min. Elution of peptides was monitored with a UV detector (215 nm, 254 nm) and a total of 36 fractions were collected, pooled into 12 fractions using a post-concatenation strategy as previously described [49] and dried under vacuum. Dried peptides were resuspended in 0.1% aqueous formic acid and subjected to LC–MS/MS analysis using a Q Exactive HF Mass Spectrometer fitted with an EASY-nLC (both Thermo Fisher Scientific) and a custom-made column heater set to 60 °C. Peptides were resolved using a RP-HPLC column (75 μm × 30 cm) packed in-house with C18 resin (ReproSil-Pur C18–AQ, 1.9 μm resin; Dr. Maisch GmbH) at a flow rate of 0.2 µl/min. The following gradient was used for peptide separation: from 5% B to 15% B over 19 min to 30% B over 80 min to 45% B over 21 min to 95% B over 2 min followed by 18 min at 95% B. Buffer A was 0.1% formic acid in water and buffer B was 80% acetonitrile, 0.1% formic acid in water. The Q Exactive HF mass spectrometer was operated in DDA mode with a total cycle time of approximately 1 second. Each MS1 scan was followed by high-collision-dissociation (HCD) of the 10 most abundant precursor ions with dynamic exclusion set to 30 seconds. For MS1, 3e6 ions were accumulated in the Orbitrap over a maximum time of 100 ms and scanned at a resolution of 120,000 FWHM (at 200 m/z). MS2 scans were acquired at a target setting of 1e5 ions, maximum accumulation time of 100 ms and a resolution of 30,000 FWHM (at 200 m/z). Singly charged ions and ions with unassigned charge state were excluded from triggering MS2 events. The normalized collision energy was set to 35%, the mass isolation window was set to 1.1 m/z and one microscan was acquired for each spectrum. The acquired raw-files were converted to the mascot generic file (mgf) format using the msconvert tool (part of ProteoWizard, version 3.0.4624 (2013-6-3)) and searched using MASCOT against a murine database (consisting of 34026 forward and reverse protein sequences downloaded from Uniprot on 20190129), the six calibration mix proteins [48] and 392 commonly observed contaminants. The precursor ion tolerance was set to 10 ppm and fragment ion tolerance was set to 0.02 Da. The search criteria were set as follows: full tryptic specificity was required (cleavage after lysine or arginine residues unless followed by proline), 3 missed cleavages were allowed, carbamidomethylation (C) and TMT6plex (K and peptide N-terminus) were set as fixed modification and oxidation (M) as a variable modification. Next, the database search results were imported into the Scaffold Q+ software (version 4.3.2, Proteome Software Inc., Portland, OR) and the protein false identification rate was set to 1% based on the number of decoy hits. Proteins that contained similar peptides and could not be differentiated based on MS/MS analysis alone were grouped parsimoniously. Proteins sharing significant peptide evidence were grouped into clusters. Acquired reporter ion intensities in the experiments were employed for automated quantification and statistical analysis using a modified version of our in-house developed SafeQuant R script[48]. This analysis included adjustment of reporter ion intensities, global data normalization by equalizing the total reporter ion intensity across all channels, summation of reporter ion intensities per protein and channel, calculation of protein abundance ratios and testing for differential abundance using empirical Bayes moderated t-statistics. To meet additional assumptions (normality and homoscedasticity) underlying the use of linear regression models and t-Tests, MS-intensity signals were transformed from the linear to the log-scale. All proteins detected are presented in supplementary table 2. Finally, significantly deregulated proteins were defined as log2 (fold change) > 0.5 or log2(fold change) < −0.5, p-value < 0.01.

### Phosphoproteomics

Peptide samples were enriched for phosphorylated peptides using Fe(III)-IMAC cartridges on an AssayMAP Bravo platform as described [50]. Unmodified peptides (“flowthrough”) were subsequently used for TMT analysis. Phospho-enriched peptides were resuspended in 0.1% aqueous formic acid and subjected to LC– MS/MS analysis using an Orbitrap Fusion Lumos Mass Spectrometer fitted with an EASY-nLC 1200 (both Thermo Fisher Scientific) and a custom-made column heater set to 60 °C. Peptides were resolved using a RP-HPLC column (75 μm × 37 cm) packed in-house with C18 resin (ReproSil-Pur C18–AQ, 1.9 μm resin; Dr. Maisch GmbH) at a flow rate of 0.2 µl/min. The following gradient was used for peptide separation: from 5% B to 8% B over 5 min to 20% B over 45 min to 25% B over 15 min to 30% B over 10 min to 35% B over 7 min to 42% B over 5 min to 50% B over 3min to 95% B over 2 min followed by 18 min at 95% B. Buffer A was 0.1% formic acid in water and buffer B was 80% acetonitrile, 0.1% formic acid in water. The Orbitrap Fusion Lumos mass spectrometer was operated in DDA mode with a cycle time of 3 seconds between master scans. Each master scan was acquired in the Orbitrap at a resolution of 120,000 FWHM (at 200 m/z) and a scan range from 375 to 1600 m/z followed by MS2 scans of the most intense precursors in the Orbitrap at a resolution of 30,000 FWHM (at 200 m/z) with isolation width of the quadrupole set to 1.4 m/z. Maximum ion injection time was set to 50 ms (MS1) and 54 ms (MS2) with an AGC target set to 1e6 and 5e4, respectively. Only peptides with charge state 2 – 5 were included in the analysis. Monoisotopic precursor selection (MIPS) was set to Peptide, and the Intensity Threshold was set to 2.5e4. Peptides were fragmented by HCD (Higher-energy collisional dissociation) with collision energy set to 30%, and one microscan was acquired for each spectrum. The dynamic exclusion duration was set to 30 seconds. The acquired raw-files were imported into the Progenesis QI software (v2.0, Nonlinear Dynamics Limited), which was used to extract peptide precursor ion intensities across all samples applying the default parameters. The generated mgf-file was searched using MASCOT against a murine database (consisting of 34026 forward and reverse protein sequences downloaded from Uniprot on 2019-01-29) and 392 commonly observed contaminants using the following search criteria: full tryptic specificity was required (cleavage after lysine or arginine residues, unless followed by proline); 3 missed cleavages were allowed; carbamidomethylation (C) was set as fixed modification; oxidation (M) and phosphorylation (STY) were applied as variable modifications; mass tolerance of 10 ppm (precursor) and 0.02 Da (fragments). The database search results were filtered using the ion score to set the false discovery rate (FDR) to 1% on the peptide and protein level, respectively, based on the number of reverse protein sequence hits in the datasets. Exported peptide intensities were normalized based on the protein regulations observed in the corresponding TMT experiment in order to account for changes in protein abundance. Only peptides corresponding to proteins, which were regulated significantly with a p value ≤ 1% in the TMT analysis were normalized. Quantitative analysis results from label-free quantification were processed using the SafeQuant R package v.2.3.2 [48] (https://github.com/eahrne/SafeQuant/) to obtain peptide relative abundances. This analysis included global data normalization by equalizing the total peak/reporter areas across all LC-MS runs, data imputation using the knn algorithm, summation of peak areas per peptide and LC-MS/MS run, followed by calculation of peptide abundance ratios. Only isoform specific peptide ion signals were considered for quantification. The summarized peptide expression values were used for statistical testing of between condition differentially abundant peptides. Here, empirical Bayes moderated t-Tests were applied, as implemented in the R/Bioconductor limma package were used. All LC-MS analysis runs were acquired from independent biological samples. To meet additional assumptions (normality and homoscedasticity) underlying the use of linear regression models and t-Tests, MS-intensity signals were transformed from the linear to the log-scale. All proteins detected are presented in supplementary table 2. Finally, significantly deregulated were selected by a calculated p-value < 0.01.

### Statistical analysis for phosphoprotemics

Using the software Perseus [51], we performed PCA after Lg2 transformation and subsequent normalization by median subtraction. For the volcano-plot, we used the double sided t-test with 250 randomizations. We set the false discovery rate (FDR) to 0.05 and the S0 value to 0.1 as threshold. To generate hierarchical clustering with z-scored data, we used Spearman correlation for the distance of both trees, leaving the other parameters as default.

### Pathway enrichment analysis

Pathway enrichment analysis for proteins with altered phosphorylation sites (p-value < 0.01) between iAdRiKO and control mice was performed by the Database for Annotation, Visualization and Integrated discovery (DAVID) v6.8 [52] (https://david.ncifcrf.gov/). Settings for DAVID were default. The background list consisted of all proteins detected in the phosphoproteome data set (supplementary table 2). The full analysis can be found in supplementary table 3.

### Immunofluorescence staining of WAT sections

Inguinal WATs were fixed overnight in 4% formalin at room temperature, dehydrated, embedded in paraffin, and cut into 5μm thick sections. For immunofluorescence staining, sections were rehydrated and antigen retrieved by boiling sections in target retrieval Solution (Dako) for 20 min in a KOS Microwave HistoSTATION (Milestone Medical). Sections were blocked using Protein block serum-free ready-to-use (Dako) and then incubated in primary antibody diluted in antibody diluent with background reducing component (Dako) overnight at 4°C. Primary antibodies against tyrosine hydroxylase (1:500, Millipore, Cat#AB1542), CGRP (1:200, Enzo, BML-CA1137-0100), GAP43-pS41 (1:200, R&D system, PPS006), Neurofilament heavy polypeptide (1:1000, Abcam, ab8135), Synaptophysin (1:200, Abcam, ab32127) and Adiponectin (1:500, Invitrogen, PA1-054) were used. After washing, sections were incubated in secondary antibody (1:500) in antibody diluent with background reducing component (Dako) for 1 hour at room temperature. Secondary antibodies against rabbit (Invitrogen, A21070 or A21443), sheep (Invitrogen, A21436) and mouse (Invitrogen, A11004) were used. Finally, sections were stained with NucBlue Live Cell Staining ReadyProbes reagent (Invitrogen, R37605) for 2 min and mounted with VECTASHIELD (Vector, H-1000). Images were obtained using Applied Precision Deltavision CORE system (Leica) and analyzed with OMERO [53]. For quantification, maximal intensity levels were used.

### Whole mount imaging using inguinal WAT depots

Tissues were harvested from mice after intracardiac perfusion with 4% Paraformaldehyde (PFA) and further fixed in 4% PFA overnight at 4 °C. For immunolabeling, tissue clearing and high-resolution volumetric imaging, we developed a modified protocol with the focus on tissue structure and fluorescence preservation. First steps are described by Jingyi Chi et al. 2018 [32]. In short, tissues were washed with PBS, dehydrated using a methanol/B1n buffer (0.3 M Glycine, 0.1% (v/v) Triton-X, pH 7), delipidated using dichloromethane, bleached overnight with 5% H2O2 at 4°C and rehydrated in a reversed methanol/B1n buffer series. For the subsequent immunolabeling tissues were incubated with primary antibodies against TH (1:500, Millipore, Cat#AB1542) and CGRP (1:750, ImmunoSTAR, Cat#24112) diluted in PTxwH buffer (PBS, 0.1% Triton-X (v/v), 0.05% Tween20 (v/v), 2 µg/ml Heparin) and secondary antibodies against sheep (1:500, Invitrogen, A21436 or A21448) and/or rabbit (1:500, Invitrogen, A21070 or A21443). Further tissue clearing was performed using the water-based clearing method Cubic L [34]. Briefly, samples cleared by CUBIC L solution (10% N-butyldiethanolamine (v/v), 10% Triton X-100 (v/v) in ddH_2_0) followed by 2% agarose embedding and refractive index (RI) matched in CUBIC RA solution (45% antipyrine, 30% N-methylnicotinamide) before imaging. After RI matching Tissue was imaged with the Zen black software (ZEISS) in Cubic RA solution with the RI of 1.51 on a Carl ZEISS lightsheet 7 microscope equipped with the Clr Plan-Neofluar 20x/1.0 detection objective and dual side illuminated with 10x/0.2 objectives. Acquired tiles were loaded in ArivisVision4D (Arivis) for stitching, visualization, and analysis.

### CGRP ELISA

Tissues were lysed in PBS using lysis matrix D tubes (MP Biomedicals). CGRP levels in lysates were determined using mouse calcitonin gene related peptide (CGRP) ELISA kit (CSB-EQ027706MO, Cusabio) according to manufacturer’s instructions.

### Denervation surgery

Inguinal fad depots were denervated as described previously [54] in eight-week-old mice. For surgical denervation, mice were anaesthetized and incisions were made dorsally on the flank. Nerves innervating inguinal WAT were identified using a dissection microscope, cut several times and removed. For sham operation, mice were anaesthetized and incisions were made dorsally on the flank. Mice were allowed to recover after surgery and were sacrificed four weeks after surgery.

### Statistics

Sample size was chosen according to our previous studies and published reports in which similar experimental procedures were described. The investigators were not blinded to the treatment groups. All data are shown as the mean ± SD. Sample numbers are indicated in each figure legend. For mouse experiments, *n* represents the number of animals, and for imaging, *N* indicates the number of images acquired per experiment. To determine the statistical significance between two groups, an unpaired two-tailed Student’s t-test was performed. For ITT, two-way ANOVA was performed. All statistical analyses were performed using GraphPad Prism 9 (GraphPad Software, San Diego, California). A *p* value of less than 0.05 was considered statistically significant.

## Supplementary Figure and Table Legends

**Supplementary Figure 1.**
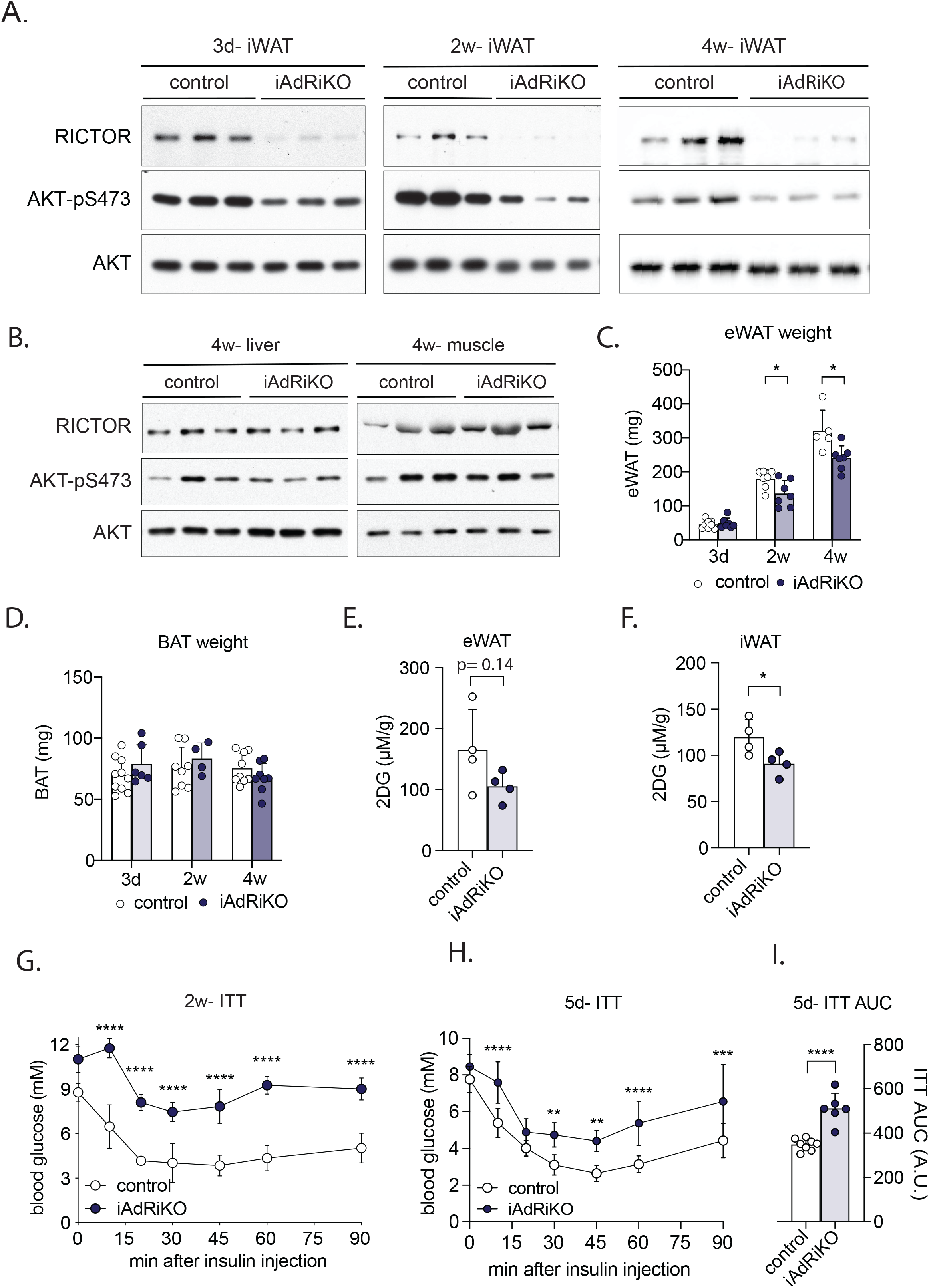
(A) Immunoblot analysis of inguinal WAT (iWAT) from control and iAdRiKO mice three days, two, and four weeks after tamoxifen treatment (n=6;6). (B) Immunoblot analysis of liver and muscle tissue from control and iAdRiKO mice four weeks after tamoxifen treatment (n=6;6). (C) Tissue weight of epididymal WAT (eWAT) in *ad libidum*-fed control and iAdRiKO mice three days (n=8;7), two (n=7), and four weeks (n=5;7) after tamoxifen treatment. Student’s t test, *p<0.05. (D) Tissue weight of brown adipose tissue (BAT) in *ad libidum*-fed control and iAdRiKO mice at three days (n=10;6), two (n=8;4) and four (n=9;8) weeks after tamoxifen treatment. (E) 2-deoxyglucose (2DG) uptake in eWAT of control and iAdRiKO mice three days after tamoxifen treatment (n=4). (F) 2DG uptake in iWAT of control and iAdRiKO mice three days after tamoxifen treatment (n=4). Student’s t-test, *p<0.05. (G) Insulin tolerance test (ITT) on control and iAdRiKO mice two weeks after tamoxifen treatment (n=4). 2-way ANOVA, **** p<0.0001. (H) ITT on control and iAdRiKO mice five days after tamoxifen treatment (n=8;6). 2-way ANOVA, **p<0.01, ***p<0.001, **** p<0.0001. (I) Area under the curve (AUC) for ITT on control and iAdRiKO mice five days after tamoxifen treatment (n=8;6). Student’s t-test, **** p<0.0001.

**Supplementary Figure 2.**
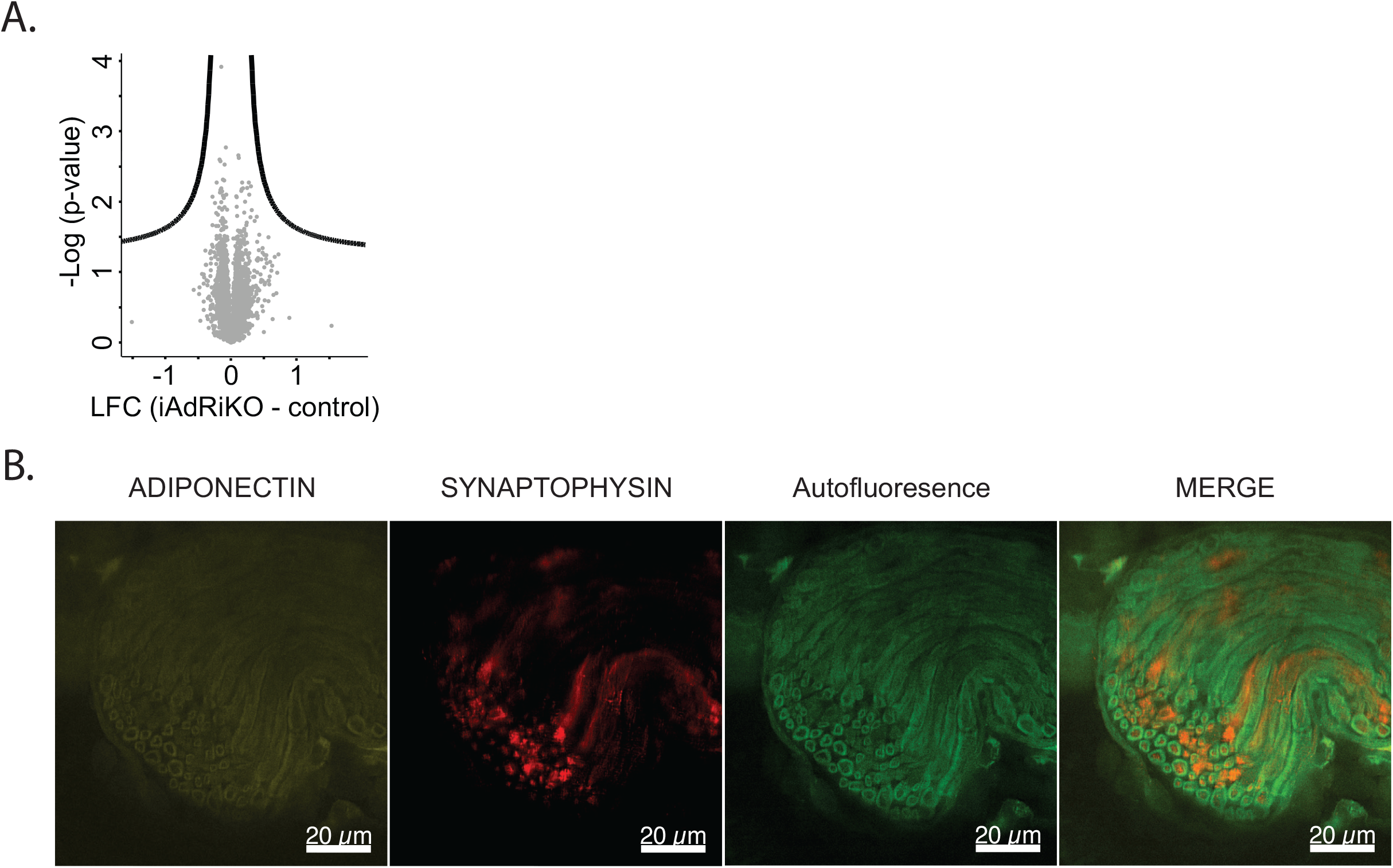
(A) Volcano-plot displaying the comparison of proteins derived from inguinal WAT (iWAT) of control and iAdRiKO mice at three days after tamoxifen treatment (n=5). LFC= Log_2_ fold change. (B) Representative image of large nerve bundles in control iWAT immunostained with both ADIPONECTIN and SYNAOTIPHYSIN (N=4). Scale bar= 20 µm.

**Supplementary Figure 3.**
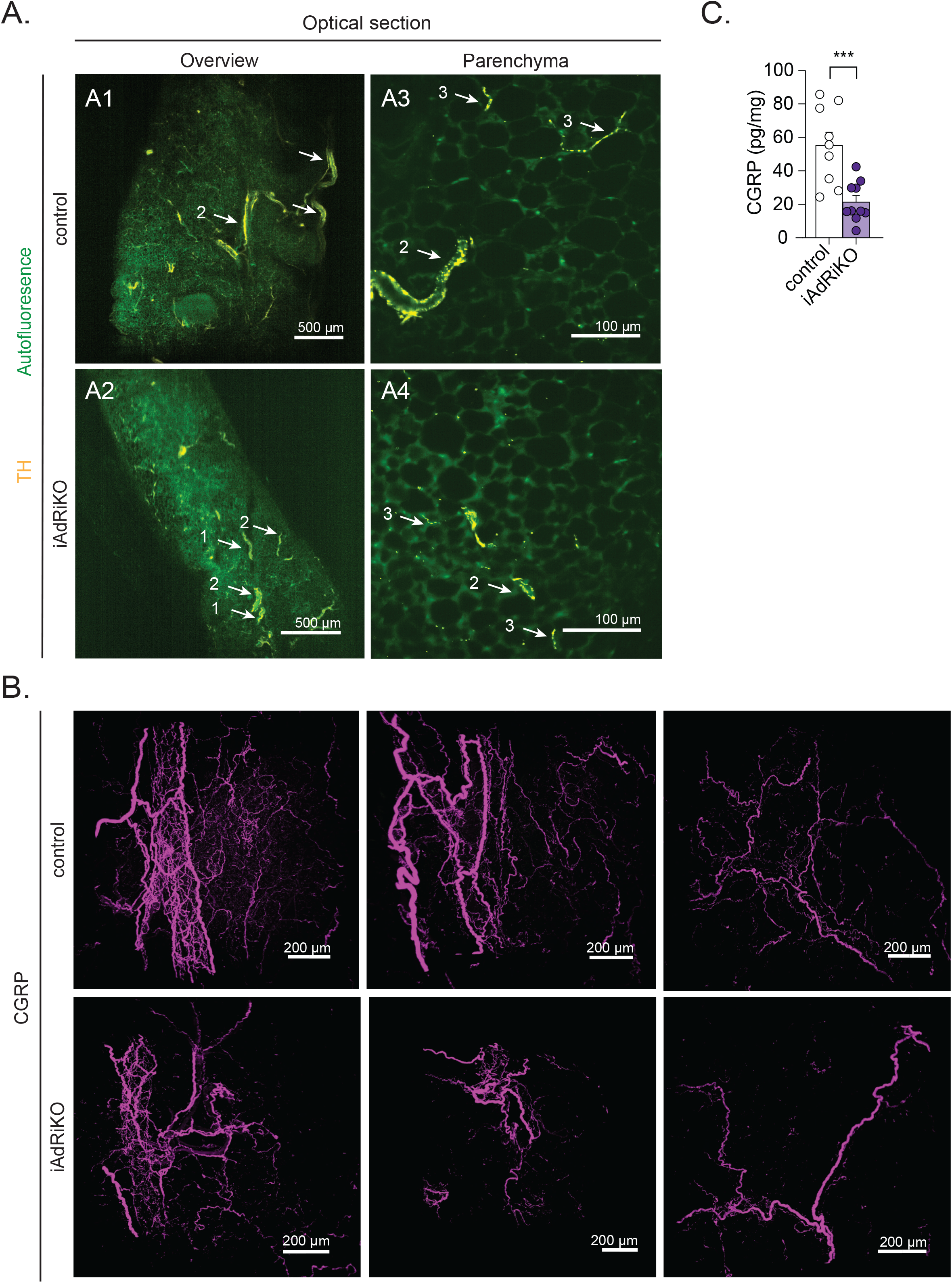
(A) Crosssections of inguinal WAT (iWAT) four weeks after tamoxifen treatment immunostained with tyrosine hydroxylase (TH; yellow). (A1-2) Low magnification cross sections of sympathetic neuronal network in control and iAdRiKO mice (N=**4;5**). Nerve bundle (1), innervation around blood vessel (2), tissue autofluorescence (green). Scale bar= 500 µm. (A3-4) High magnification cross section of sympathetic neurons in iWAT parenchyma of control and iAdRiKO mice (N=19;10). Scale bar= 100 µm. (B) High magnification projections of calcitonin gene-related peptide (CGRP)-positive sensory neurons (magenta) in iWAT of different control and iAdRiKO mice. Scale bar= 200 µm. (N=13;8, n=3;3). (C) Enzyme-linked immunosorbent assay (ELISA) for CGRP levels in iWAT of control and iAdRiKO mice (n=9). Student’s t-test, ***p<0.001.

**Supplementary Figure 4.**
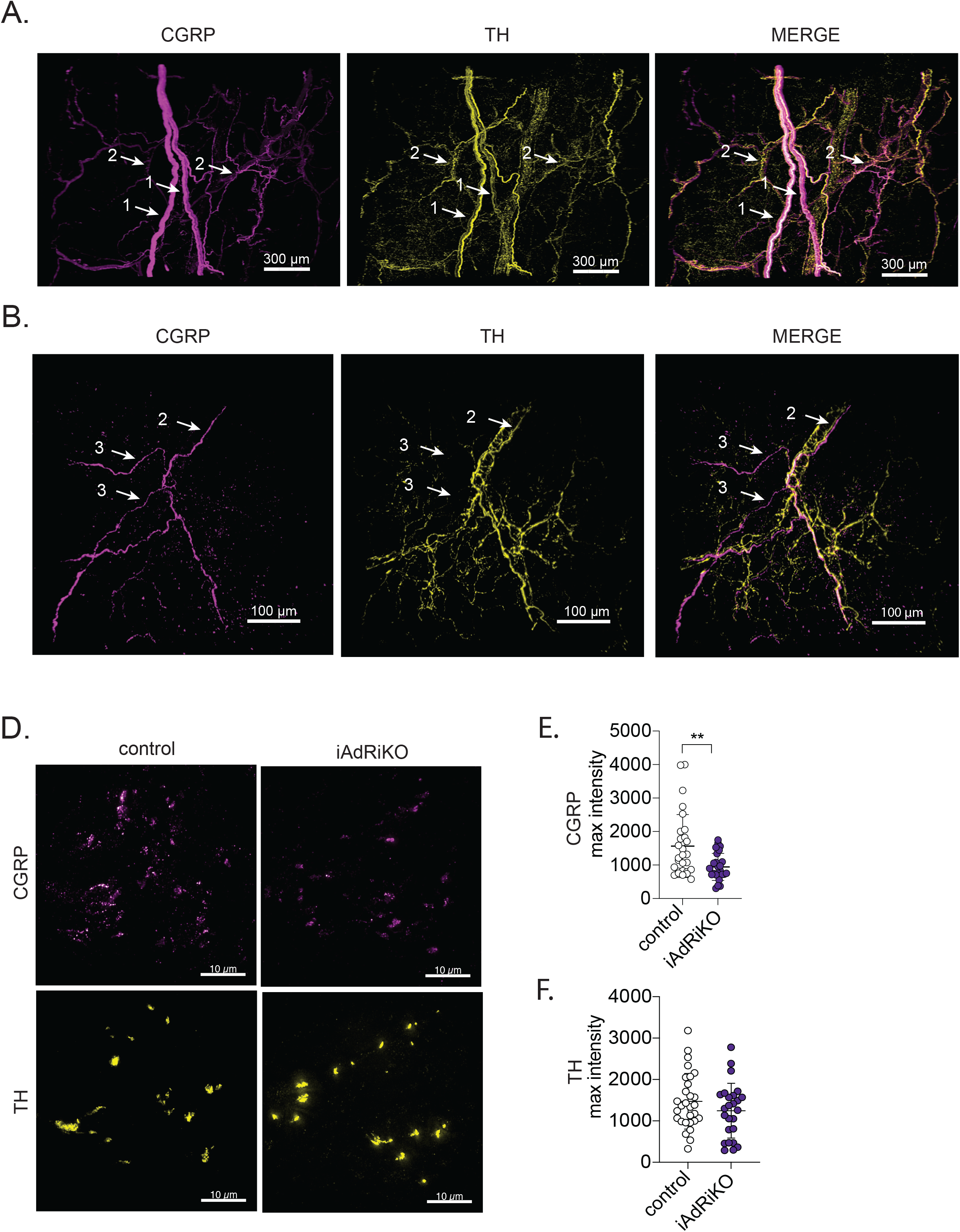
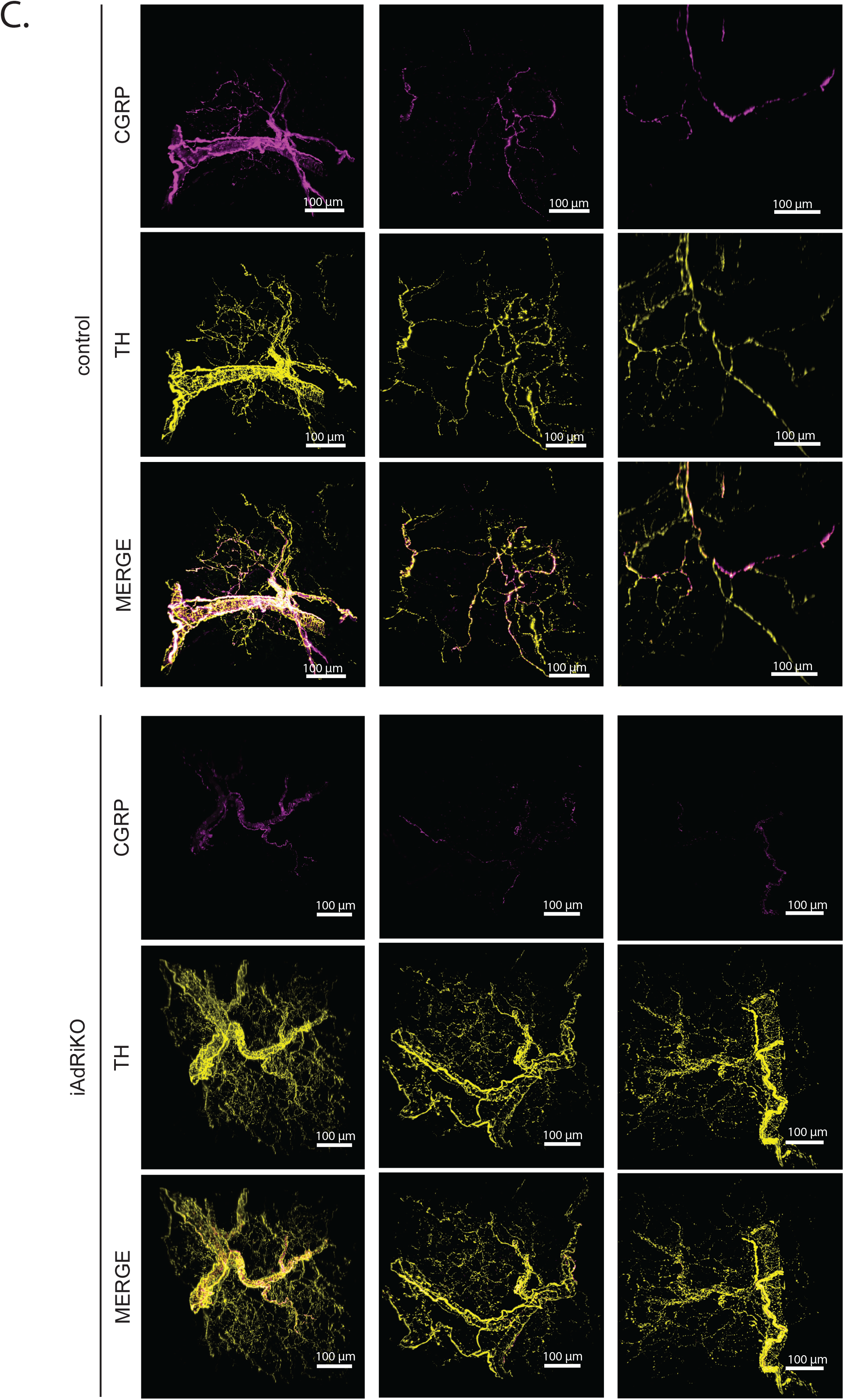
(A) 2D representative of a 3D reconstruction of an inguinal WAT (iWAT) section immunostained with calcitonin gene-related peptide (CGRP, magenta) and the tyrosine hydroxylase (TH, yellow) that illustrates how a large nerve bundle subdivides into smaller nerve bundles which in turn penetrate the tissue alongside blood vessels. Nerve bundle (1), innervation along blood vessel (2). Scale bar= 300 µm. (B) 2D representative of a 3D reconstruction of the iWAT parenchyma of both CGRP-positive (magenta) and the TH-positive (yellow) neuronal networks in iWAT. Innervation along blood vessel (2), parenchymal innervation (3). Scale bar= 100 µm. (C) 2D representative of a 3D reconstruction of inguinal WAT sections from control and iAdRiKO mice immunostained with calcitonin gene-related peptide (CGRP; magenta) and tyrosine hydroxylase (TH; yellow) (N=21;22). Scale bar= 100 µm. (D) Representative image of a large nerve bundle in control and iAdRiKO iWAT paraffine sections immunostained with CGRP (magenta) and tyrosine hydroxylase (TH, yellow) (N= 29;25). Scale bar= 10 µm. (E) Quantification of maximal intensity of CGRP-immunostaining in large nerve bundles of iWAT of control and iAdRiKO mice (N= 29;25). Student’s t-test, **p<0.01. (F) Quantification of maximal intensity of TH-immunostaining in large nerve bundles of iWAT of control and iAdRiKO mice (N= 29;25). Student’s t-test, **p<0.01.

**Supplementary Table 1**.

(A) Proteome data set from inguinal WAT (iWAT) of iAdRiKO and control mice three days after tamoxifen treatment

(B) Phosphoproteomics data set from iWAT of iAdRiKO and control mice three days after tamoxifen treatment. Significant hits are highlighted in yellow (p< 0.01).

**Movie 1**

Calcitonin gene-related peptide (CGRP)-positive neuronal network (magenta) in the inguinal WAT parenchyma of a control mouse

**Movie 2**

CGRP-positive neuronal network (magenta) in the inguinal WAT parenchyma of an iAdRiKO mouse

**Movie 3**

Overview of both CGRP- (magenta) and tyrosine hydroxylase (TH; yellow)- positive neuronal networks in inguinal WAT of a control mouse.

**Movie 4**

CGRP- (magenta) and TH- (yellow) positive neuronal networks in the inguinal WAT parenchyma of a control mouse.

**Movie 5**

CGRP- (magenta) and TH- (yellow) positive neuronal networks in the inguinal WAT parenchyma of an iAdRiKO mouse.

